# Evaluating Sample-Size Efficiency and Sensitivity of Tractometry in Alzheimer’s Disease

**DOI:** 10.1101/2025.11.11.687878

**Authors:** Yixue Feng, Julio E. Villalón-Reina, Iyad Ba Gari, Jonathan Davis Alibrando, Talia M. Nir, Neda Jahanshad, Bramsh Q. Chandio, Paul M. Thompson

## Abstract

Tractometry allows quantitative analysis of white matter microstructure along the brain’s fiber tracts, but the impact of study design parameters—such as sample size and along-tract resolution—on sensitivity and specificity is not well understood. In this study, we conducted tractometry bootstrap analysis using linear-mixed models across four diffusion tensor imaging (DTI) metrics to systematically evaluate how these factors affect the detection of dementia- and amyloid-related effects. While coarser along-tract segments yield greater sensitivity and higher mean effect sizes, finer segments tend to produce higher peak effect sizes, revealing more spatially localized effects. Dementia-related effects were more widespread and detectable with fewer subjects, whereas amyloid-related effects were more subtle and localized, requiring larger cohorts to detect them. These findings highlight that tractometry offers improved spatial specificity and can reliably detect small, fine-scale effects, but study design should be tailored to specific research questions, considering the expected spatial extent and magnitude of effects, to optimize sample size efficiency and interpretability.

## 1. INTRODUCTION

By sampling diffusion MRI (dMRI) derived measures along the trajectories of major white matter (WM) bundles reconstructed using tractography, tractometry [1] has become a central approach for characterizing microstructural alterations, and has been used to study Alzheimer’s disease, as well as psychiatric and developmental conditions [2, 3, 4]. Compared to voxel-based methods such as Tract-Based Spatial Statistics (TBSS) [5], or methods where diffusion measures are averaged over large WM regions of interest (ROIs), tractometry provides higher spatial resolution by mapping white matter parameters at a finer anatomical scale. However, these high-dimensional analyses introduce unique challenges for study design and statistical inference: splitting up bundles into multiple correlated segments can produce hundreds to thousands of statistical tests across the brain. Controlling the false discovery rate (FDR)[6] while maintaining sensitivity to biologically meaningful effects, therefore, will yield maps of effects that depend on sample size, along-tract resolution (i.e,. number of segments along tracts/segment length), and on the magnitude and spatial extent of the underlying disease effects. Most tractometry approaches, by default, divide tracts into 100 nodes/segments along the length of the tracts[7, 8, 9]. In this study, for the first time, we assessed how these parameters influence the detection power in tractometry analyses using diffusion MRI data from the Health & Aging Brain Study (HABS-HD) [10] cohort to study the effect of dementia and amyloid burden on white matter microstructure, mapped throughout the brain in 3D. Using repeated bootstrap sampling, we set out to assess how the proportion of bundle segments that survive a global FDR correction varies in a way that depends on the available sample size and segment lengths. By comparing diffusion metrics and along-tract sampling resolution, we identified combinations that yielded the most efficient detection of disease-related changes and provide practical guidance on optimizing future tractometry studies for statistical power and spatial specificity.

## 2. METHODS

### 2.1. Data & Preprocessing

We analyzed dMRI from 3,049 participants in the HABS-HD dataset [10](1,912 F/1,1137 M, mean age 65.55±8.70), collected on two scanners (1,724 Skyra/1,325 Vida). The dataset included 2,238 cognitively normal (CN) participants, 612 with mild cognitive impairment (MCI), and 199 with dementia. Using amyloid-sensitive positron emission tomography (PET), global brain amyloid-*β* burden was derived from 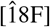 florbetaben SUVR values (FBB CL) normalized by the whole cerebellum as the reference region and converted to centiloid units for 2,044 participants (1,487 CN/434 MCI/123 with Dementia). The centiloid scale standardizes amyloid PET measurements, where 0 represents the mean signal in young amyloid-negative controls and higher values represents that of typical Alzheimer’s patients, enabling direct comparison across tracers and analysis methods[11]. Details of dMRI preprocessing may be found in [3]. We generated 87 WM bundles using a bundle tractography method [12] with a modified version of the population-averaged HCP1065 Young Adult atlas [13] as reference. We discarded 3 bundles (the left parahippocampal parietal cingulum and bilateral medial forebrain bundles) as they had less than 30% of subjects with successful reconstructions. We used BUndle ANalytics (BUAN) [9] to derive along-tract microstructural profiles using a weighted mean approach[14], where for each subject’s bundle, each point on a streamline is assigned to a segment based on the closest centroid from the atlas bundle, and DTI measures projected onto these points are weighted by their distance to the atlas bundle centroid and averaged within each segment. For each bundle, we computed along-tract profiles with 4 along-tract resolutions at 5-mm, 10-mm, 25-mm and 50-mm segment length (*l*), where the number of segments (*S*_*l*_) was calculated as the average streamline length of the atlas bundle divided by *l*. Bundles shorter than 50-mm remain as one segment. The total number of segments across all 84 tracts (*S*) for each *l* were 1619, 791, 310, and 150, respectively.

### 2.2. Tractometry Bootstrapping

In our first analysis, we assessed diagnostic group (DX) differences in 4 DTI (DTI) metrics —FA, MD, RD, and AxD —using a linear mixed-effects model (LMM) fit at each bundle segment. These are all measures that show associations with dementia in both along-tract and region of interest analyses[15, 2]. CN participants were labeled as controls, and, for the purposes of this analysis, cognitively impaired participants (those with MCI or dementia) were labeled as cases. The model was defined as

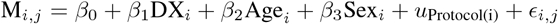

where M_*i,j*_ is the DTI metric for subject *i* at segment *j, β*_0_ is the intercept, *β*_1..3_ are the fixed effect, *u*_Protocol(i)_ is the random intercept for MRI protocol, *ϵ*_*i,j*_ is the residual error. In the second analysis, we modeled DTI metrics as a function of amyloid burden, quantified by Centiloid (FBB CL), among CN participants (N=1487):

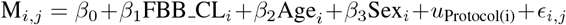

For both analyses, bootstrap resampling was used to evaluate the stability and sensitivity of along-tract effects across varying experiment conditions. Specifically, for each sample size *S* and segment length *l*, subjects were sampled with replacement for 500 bootstrap iterations. In the case-control analysis, *S* subjects per group were sampled, where *S* = 100, 200, …, 800. In the amyloid analysis, *S* CN subjects with available amyloid measures are sampled where *S* = 100, 200, …, 1200. After fitting the LMM, multiple comparisons across all segments and bundles were corrected using the Benjamini–Hochberg FDR procedure (*q <* 0.05)[6].

In this procedure, all segment-level *p*-values across bundles are first sorted in ascending order:

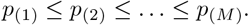

For a target false discovery rate *q* = 0.05, the Benjamini– Hochberg threshold is defined as the largest *p*_(*k*)_ satisfying

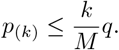

The *global FDR proportion* was then defined as the fraction of all segments with *p*_*i*_ ≤ *p*_(*k*)_, representing the proportion of bundle segments surviving global FDR correction.

The proportion of bundle segments surviving global FDR correction, or *global FDR proportion*, was computed for each bootstrap, sample size, and segment length. For each tract segment, standardized effect sizes, or *partial d*, were computed as the ratio of the fixed effect coefficient to the residual standard deviation, which quantifies the standardized magnitude associated with the predictor conditional on age, sex, and protocol. At the bundle level, we computed bundle-wise FDR proportion, the mean and max partial *d* across all segments within a bundle. These metrics summarize the spatial extent and magnitude of along-tract effects, which enables comparing sensitivity patterns across bundles. We note that bundle-level and global FDR are not always reported in tractometry publications, but we report both for completeness.

## 3. RESULTS

### 3.1. Global FDR Proportion Evaluation

To first evaluate the global sensitivity of our tractometry analyses with different sample sizes and along-tract resolutions, we plotted the global FDR proportion for both the cognitive impairment/dementia and amyloid analyses in **Fig 1** and **2**. In both analyses, the global FDR proportion increased with sample size, indicating greater statistical power to detect an effect in larger cohorts. MD and RD consistently showed higher global FDR proportions in the case-control analysis, whereas FA and RD were more sensitive to amyloid effects. Across all sample sizes, the global FDR proportion was higher in the case-control analysis, which supports literature suggesting that WM alterations associated with dementia are more widespread and pronounced [16, 17], whereas early amyloid accumulation in CN adults is more localized and has more subtle effects on WM microstructure [18].

**Fig. 1.**
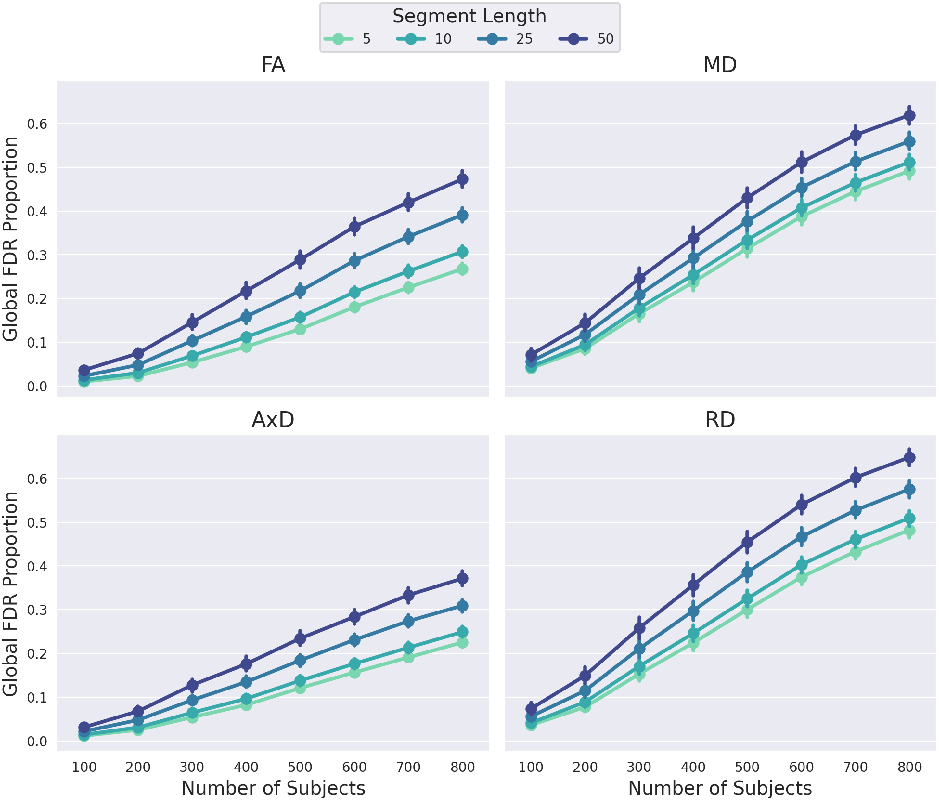
Global FDR proportion for the case-control analysis. The error bars show the 95% confidence interval across 500 bootstraps. For conditions that affect the entire brain to some extent, these curves should eventually reach 1.0, whereas effects that are truly localized would converge in theory to a fixed proportion of the tracts, below 1.0.

**Fig. 2.**
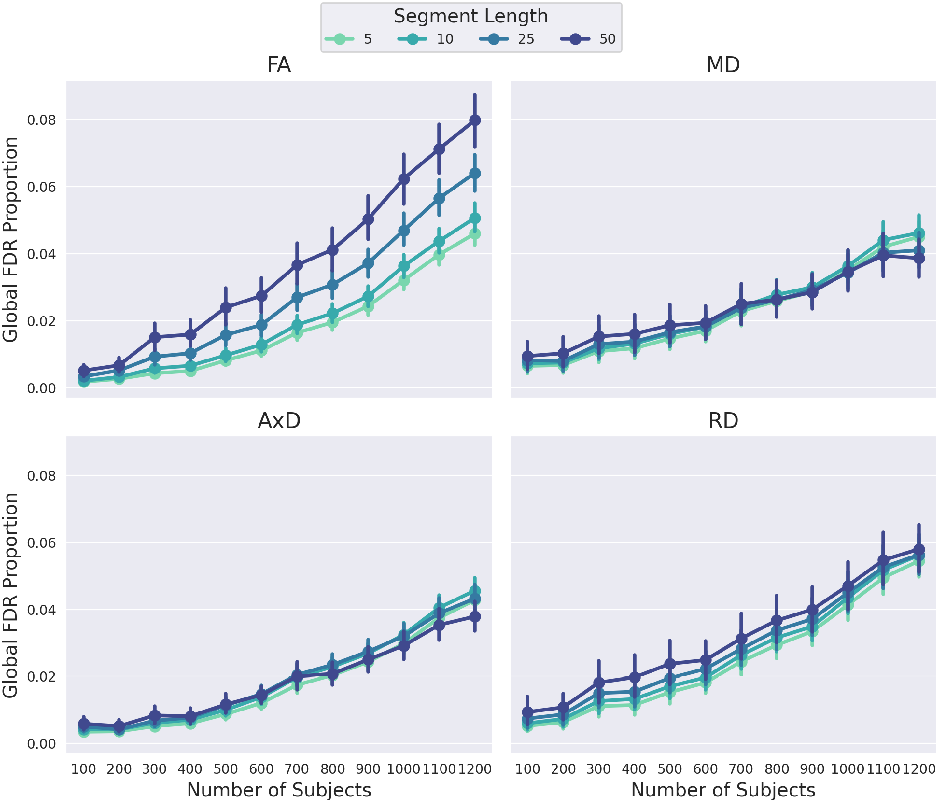
Global FDR proportion for the amyloid analysis. The error bars show the 95% confidence interval across 500 bootstraps.

We also observed an effect of along-tract resolution on global FDR proportion. In the case-control analysis, larger segment lengths generally produced higher FDR proportions across all DTI metrics, likely due to the fact that longer segments result in fewer multiple comparisons or can better capture distributed microstructural changes. In longer segments, spatially coherent effects spanning several segments are combined to improve sensitivity, as their aggregation amplifies the signal relative to noise. In contrast, in the amyloid analysis, this effect was observed primarily for FA, whereas the global FDR proportion was comparable across different choices of segment lengths for MD, RD and AxD. Interestingly, at larger sample sizes (S*>*1000), shorter segment lengths became more sensitive for MD and RD. These results suggest that in more widespread pathology, such as dementia, longer segments may better aggregate consistent microstructural changes, whereas for early or subtle WM alterations, such as amyloid accumulation, finer along-tract resolution may capture more localized effects that are otherwise averaged out in longer segments.

### 3.2. Bundle-Level Effect Size Evaluation

For each analysis, bundles were ranked based on the highest bundle FDR proportion across all DTI metrics and at the largest sample size. In the case-control analysis, the cingulum–parahippocampal bundle (CG PH) with RD showed the highest proportion, whereas in the amyloid analysis, the parietal corpus callosum body (CC Body P) with FA was most prominent. We plot their peak and mean partial *d* and bundle FDR proportion across different experiment settings in **Figure 4(a) and 3(a)**. Effect sizes stabilized with increasing sample size, as shown by the reduced variability across bootstraps. However, both the peak and mean effect sizes also showed a moderate decrease, particularly in the left CG PH, indicating that smaller sample sizes may inflate or inaccurately estimate the observed effect sizes. For the CC Body P, shorter segments consistently produced lower mean partial *d* and FDR proportion, but higher peak partial *d*. A similar pattern was observed for CG PH, except *l*=25mm, which yielded high mean partial *d* consistently and peak partial *d* at higher sample sizes (*S >* 500 per group).

**Fig. 3.**
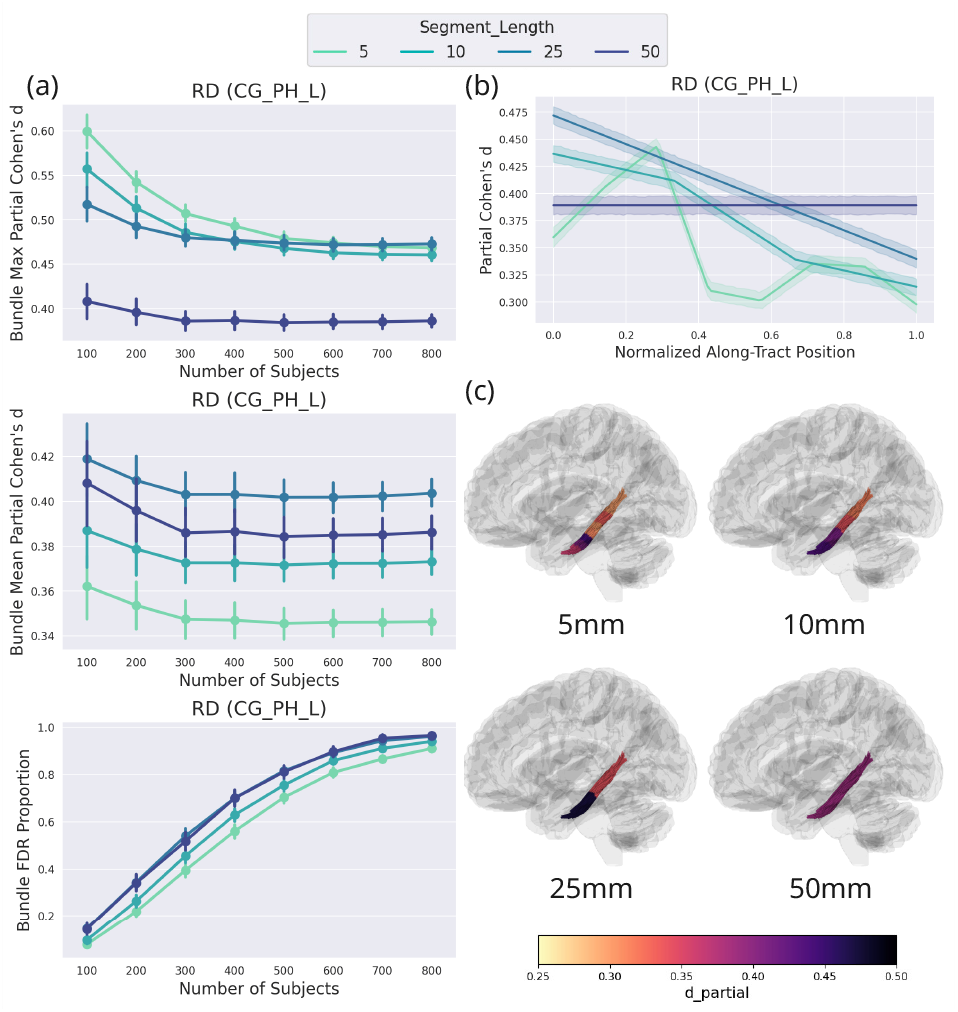
Effect size evaluation for CG PH L (the left cingulumparahippocampal bundle) in the case-control analysis. This tract is of key interest in dementia research as it connects the hippocampus and parahippocampal gyrus with the medial parietal and cingulate regions, key hubs of memory circuitry affected in Alzheimer’s disease. **(a)** Bundle FDR proportion, maximum and mean partial *d* under different experiment conditions. **(b)** Along-tract partial *d* for 4 segment lengths across bootstrap samples, also plotted on atlas bundles in **(c)**.

**Fig. 4.**
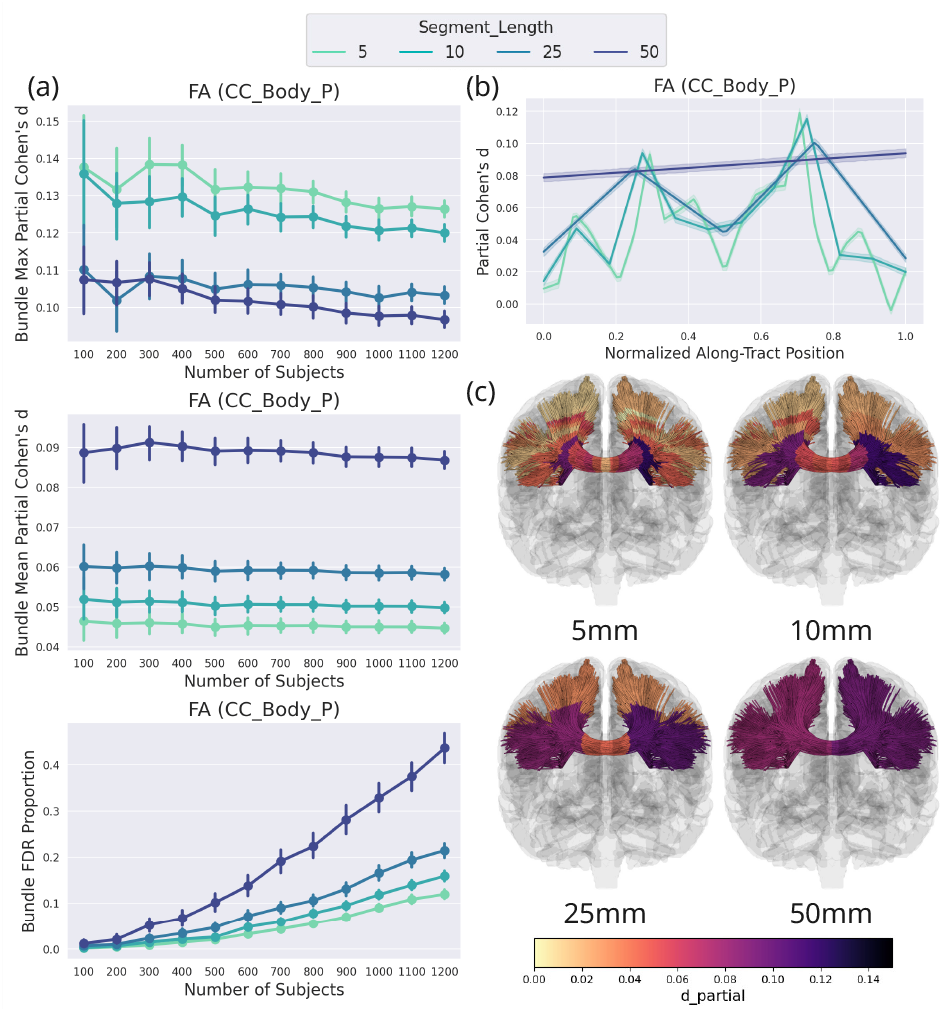
Effect size evaluation for CC Body P (the parietal section of the corpus callosum body) in the amyloid analysis. Bundle FDR proportion, max and mean partial *d* under different experimental conditions. **(b)** Along-tract partial *d* for 4 segment lengths across bootstrap samples, also plotted on atlas bundles in **(c)**.

To further illustrate these patterns, we plotted the along-tract partial *d*, averaged across bootstraps, for the largest sample size used in **Figure 4(c) and 3(c)**. Partial *d* values computed at the original segment centers were linearly interpolated to 100 evenly spaced positions, to enable direct comparison across different along-tract resolutions **Figure 4(b) and 3(b)**. For the CC Body P, high along-tract resolutions (*l*=5 and 10 mm) identified the largest effects in regions near the *corona radiata*, likely reflecting regions of crossing fibers and selective WM degeneration, consistent with prior studies using ROI-based analysis [19, 17] and tractometry [2]. When using longer segments, these focal effects appear to drive the overall bundle-level signal, resulting in an apparently broader spatial extent of the effect. This is also evident in the difference in FDR proportion across segment lengths, where the FDR proportion converges to below 0.2 for small segment lengths and but approaches above 0.4 for *l*=50mm at large sample sizes. In contrast, bundle FDR proportion for CG PH is more similar across segment lengths, approaching values above 0.9, indicating that the effect is more widespread and can be reliably detected at different along-tract resolutions. Overall, these findings suggest that while longer segments tend to capture stronger average effects across the bundle, shorter segments are more sensitive to localized peak effects.

## 4. DISCUSSION

In this study, we examined how sample size and along-tract resolution influence the sensitivity and spatial specificity of tractometry analysis. We used bootstrap LMMs across four DTI metrics to study the effect of cognitive impairment in a case-control setting and amyloid burden in CN adults. Our findings highlight that different research questions and underlying biological effects may require tailored tractometry study designs and interpretations. When studying preclinical neurodegenerative diseases where the effects are subtle and spatially variable, statistical power may be further reduced by limited sample size and multi-site variability, emphasizing the need for principled design choices. From a practical perspective, our results provide guidance for optimizing tractometry studies. When the hypothesized effect is large and spatially widespread, coarser segments (e.g., 20-30mm) may offer better sensitivity without requiring large sample sizes. This is in line with the matched filter theorem, which states that sensitivity is maximized when the analysis scale matches the spatial extent of the true signal, allowing spatially coherent ef-fects to be integrated while minimizing noise. In tractometry, spatially coherent effects across neighboring segments can be combined to boost power, analogous to smoothing in voxelwise analyses. Conversely, when the expected effect is small and localized, finer segments (5-10 mm) improve specificity and help isolate the local maxima of effect size, provided that the sample size is sufficient to support additional multiple comparisons. However, segments that are too small may introduce higher variability, especially in smaller cohorts or bundles with large diameters or fanning geometries, where voxel-level heterogeneity can obscure true effects. Sample size and along-tract resolution should therefore be considered jointly during study planning to balance sensitivity and specificity. In future work, we will further evaluate the statistical power of tractometry using simulated data to assess the relationships between sample size, detection sensitivity, and along-tract resolution. Such analyses would enable quantitative recommendations for tractometry study design, improving the efficiency and reliability of WM analysis. Adaptive sampling strategies—such as adaptive Benjamini–Hochberg procedures—can further concentrate testing in regions where signal is most likely to occur, thereby improving detection power while maintaining FDR control[20].

## 5. ACKNOWLEDGMENTS

This study was supported by the National Institutes of Health under grants RF1NS136995, R01AG087513, T32AG058507, R01MH134004 and S10OD032285 and the US Alzheimer’s Association under grant AARG-23-1149996.

